# Oribatid Mites Supply Tetrodotoxin to Poisonous Newts

**DOI:** 10.64898/2026.03.03.709434

**Authors:** Yuto Nakazawa, Satoshi Shimano, Tadachika Miyasaka, Masaatsu Adachi, Toshio Nishikawa, Kazutoshi Yoshitake, Masafumi Amano, Shigeru Sato, Kentaro Takada

## Abstract

Tetrodotoxin (TTX) is a potent neurotoxin widely distributed among marine and terrestrial animals, yet how terrestrial vertebrates acquire this toxin has been debated for decades. Here, we identify oribatid mites as the primary dietary sources of TTX for the Japanese fire-bellied newt, *Cynops pyrrhogaster*. We detect TTX and its analogs in the mites *Scheloribates processus* and *Galumna* sp. KM1, which are preferentially consumed by terrestrial newts. In a controlled toxification experiment, juvenile newts accumulated TTX after consuming TTX-bearing mites, directly demonstrating dietary sequestration. Our results demonstrate that the early terrestrial stage is critical for lifelong TTX acquisition in newts. Notably, we also detect pumiliotoxin, a poison-frog alkaloid, in *S. processus*, suggesting that newts and poison frogs have convergently evolved to exploit toxin-bearing microarthropods for chemical defense. We propose a “terrestrial toxic web” hypothesis in which oribatid mites act as a shared environmental reservoir linking putative microbial toxin production to vertebrate chemical defense, highlighting microarthropods as key mediators in terrestrial toxin flow.

---

Tetrodotoxin (TTX) is a potent neurotoxin that blocks voltage-gated sodium channels (VGSCs) in vertebrates ^1,2^. It has been found in a wide range of marine animals, such as pufferfish, blue-ringed octopuses ^3,4^, xanthid crabs ^5,6^, starfish ^7,8^, and polyclad flatworms ^9–12^, as well as in terrestrial animals such as *Atelopus* frogs ^13–15^, various newts ^1,16,17^, caddisfly larvae ^18^, and *Bipalium* flatworms ^19^. Despite almost a century of research since its initial discovery in pufferfish, the origin and biosynthetic pathway of TTX remain unresolved.

Recent studies have begun to elucidate the sequestration mechanism of TTX in marine animals. Pufferfish are attracted to 5,6,11-trideoxyTTX (TDT), a co-occurring TTX analog, suggesting a sensory mechanism for toxin acquisition ^20,21^. TDT has been identified in a variety of marine taxa, implying that pufferfish may feed on TTX-bearing organisms through chemical cues, thereby accumulating TTX through the food chain. Indeed, toxic flatworms have been implicated as a dietary source contributing to the toxification of pufferfish ^22,23^.

While the origin of marine TTX is increasingly understood through trophic interactions, its source in terrestrial TTX-bearing organisms remains unclear. In the Japanese fire-bellied newt, *Cynops pyrrhogaster*, juveniles reared on a TTX-free artificial diet gradually lose toxicity ^24^, whereas wild juveniles possess significant amounts of TTX ^24^. TTX levels in the red-spotted newt, *Notophthalmus viridescens*, disappear after six years of non-toxic feeding ^25^, supporting the idea that newts sequester TTX from external sources. Accordingly, several terrestrial organisms have been proposed as dietary sources, including caddisfly larvae ^18^ and *Bipalium* flatworms ^19^. However, the TTX in caddisfly larvae may derive from newt eggs, and predator–prey interactions between newts and *Bipalium* remain unverified. Thus, these studies provide only indirect evidence for an exogenous origin of TTX. In contrast, the rough-skinned newt, *Taricha granulosa*, can maintain or even increase TTX under long-term captivity ^26^, and skin-gland TTX levels partially recover after experimental depletion ^27^, suggesting endogenous production.

The composition of TTX analogs differs between terrestrial and marine species. TDT is widely distributed in marine species but absent in terrestrial animals, while 6-*epi*TTX and 8-*epi*TTX are terrestrial-specific analogs found exclusively in newts ^28^. Furthermore, putative biosynthetic intermediates also differ: Cep-210 and Cep-212 in terrestrial species versus Tb-210, Tb-226, Tb-242, and Tb-258 in marine species ^29,30^. These differences suggest that TTX and its analogs are likely synthesized through distinct biosynthetic pathways in each environment, terpene-like in terrestrial and polyketide-like in marine organisms ^29,30^, which are important implications for the evolution of chemical defense in TTX-bearing organisms.

To address the unresolved question of TTX origin in terrestrial vertebrates, we focused on the Japanese newt *Cynops pyrrhogaster*, which contains both TTX ^17,24,28^ and the putative intermediate Cep-210 ^29^. *C. pyrrhogaster* has a biphasic life history, transitioning from the aquatic larval stage (lasting several months) to a terrestrial phase (lasting 2–3 years) before returning to water for reproduction, after which individuals spend most of their lives in aquatic habitats. Here, we aimed to identify the dietary sources of TTX in this species by quantifying TTX levels across developmental stages, analyzing stomach contents, and screening potential prey organisms for TTX.

## Results and Discussion

### Tetrodotoxin (TTX) and Cep-210 accumulation across development in the newt

We first quantified TTX and Cep-210 in *C. pyrrhogaster* across six developmental stages: egg, larva, terrestrial juvenile, terrestrial subadult, terrestrial adult, and aquatic adult (Fig. 1A–C, Table S1). Our analysis revealed a marked increase in TTX and Cep-210 levels during the early terrestrial phase; TTX was undetectable or present only at trace levels during the egg and larval stages, but marked accumulation began at the terrestrial juvenile stage, with a significant increase from terrestrial juveniles (2.0 ± 0.23 μg) to terrestrial subadults (7.1 ± 3.1 μg) (*P* = 0.005) (Fig. 1B). Cep-210 showed a similar pattern, with a marked increase from terrestrial juveniles (0.034 ± 0.0051 μg) to terrestrial subadults (0.95 ± 0.69 μg) (*P* = 0.013) (Fig. 1C). In contrast, no significant difference was observed between terrestrial adults and aquatic adults in either TTX (12.3 ± 8.1 μg vs 12.0 ± 6.5 μg; *P* = 0.93) or Cep-210 (2.57 ± 3.71 μg vs 1.40 ± 1.64 μg; *P* = 0.463), even though aquatic-stage adults are generally older than terrestrial ones (Fig. 1B and Fig. 1C). These results are consistent with previous reports that TTX levels increase during the terrestrial stage ^24,31^ and that aquatic larvae contain little or no TTX ^31^. Our findings further demonstrate that the putative biosynthetic intermediate Cep-210 ^29^ exhibits a similar developmental pattern, increasing markedly during the terrestrial phase. In *C. pyrrhogaster*, the ecological significance of the terrestrial stage has remained poorly understood. Our results support the conclusion that TTX sequestration occurs primarily during the terrestrial stage, when newts are more exposed to terrestrial predators. The acquisition of TTX as a chemical defense early in terrestrial life may thus represent a critical adaptive strategy for survival during this vulnerable developmental period. Overall, our results highlight the terrestrial life stage as a pivotal period during which *C. pyrrhogaster* acquires TTX that provides lifelong chemical protection.

**Figure 1.**
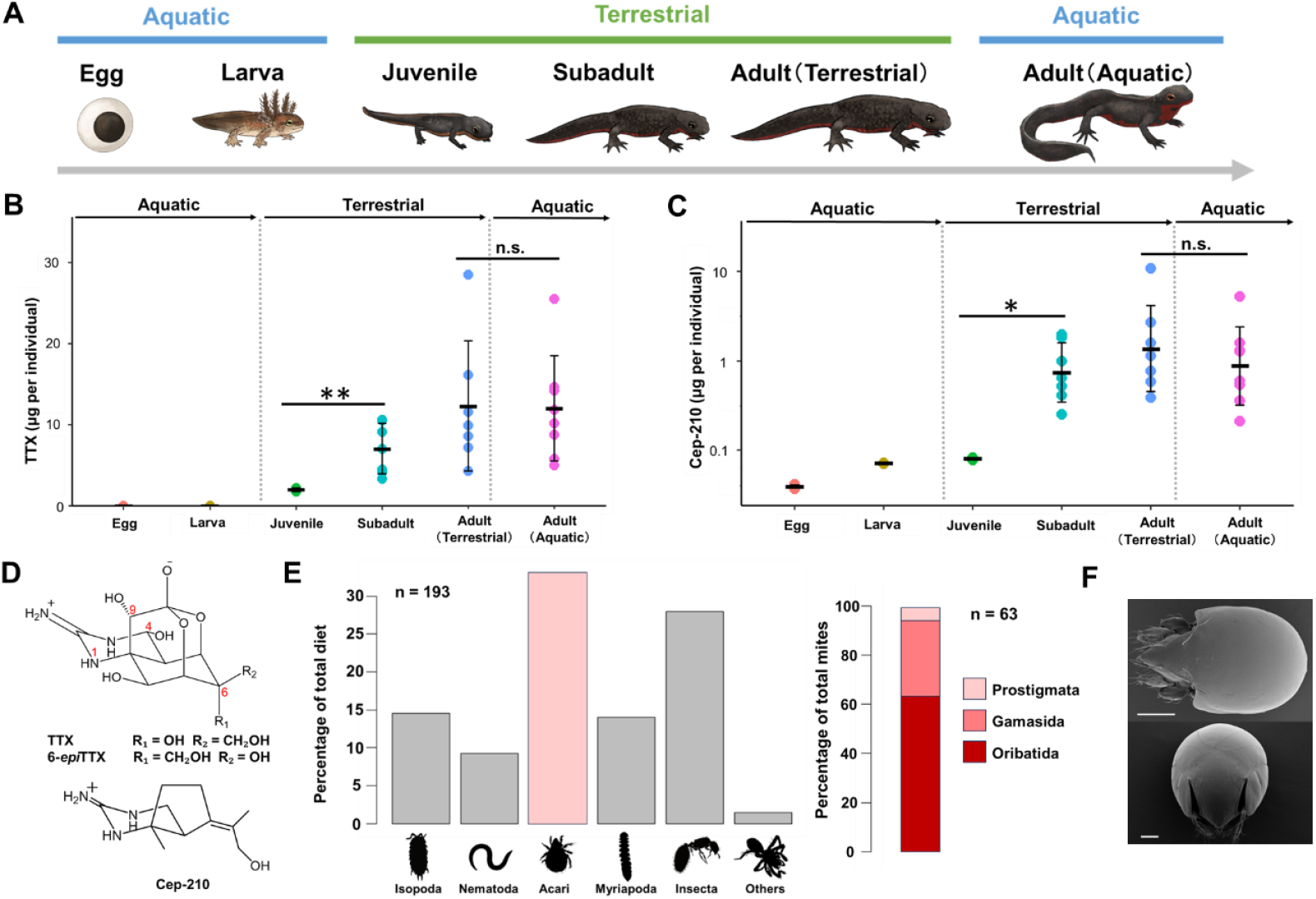
Tetrodotoxin (TTX) and Cep-210 accumulation across development and dietary composition during the terrestrial stage of the Japanese fire-bellied newt, *C. pyrrhogaster*. **A**, Schematic life cycle of *C. pyrrhogaster*. Eggs are laid in water and hatch into aquatic larvae with external gills. After several months, larvae metamorphose into juveniles that enter a terrestrial stage. Juveniles grow on land and, after reaching maturity (Adult, Terrestrial), return to water to breed (Adult, Aquatic). **B**, TTX levels across six developmental stages. **C**, Cep-210 levels across six developmental stages. Sample sizes are identical in **B** and **C**: egg (n = 6), larva (n = 4), terrestrial juvenile (n = 3), terrestrial subadult (n = 7), terrestrial adult (n = 7), and aquatic adult (n = 8). Horizontal bars and error bars indicate mean ± SD; \**P* < 0.05; \*\**P* < 0.01; n.s., not significant. **D**, Chemical structures of TTX and 6-*epi*TTX (top) and Cep-210 (bottom). **E**, Dietary composition of 193 prey items from the stomach contents of 10 terrestrial newts (left), and taxonomic composition of 63 mites (right). **F**, SEM images of the two frequently detected oribatid mite species in newt stomach contents: *Scheloribates processus* (top) and *Galumna* sp. KM1 (bottom). Scale bars, 100 μm.

### Stomach Contents Analysis

We analyzed stomach contents in ten terrestrial newts and identified a variety of small soil invertebrates, totaling 193 prey items, including Isopoda (14.5%), Nematoda (9.3%), Acari (32.6%), Myriapoda (14.0%), and Insecta (28.0%) (Fig. 1E, Table S2). Consistent with previous reports indicating that terrestrial newts preferentially feed on soil mites ^32^, we further classified 63 mite individuals into the three taxonomic groups: Prostigmata (n = 4), Gamasida (n = 19), and Oribatida (n = 40) (Fig. 1E, Table S2). Oribatid mites were the most abundant prey group in *C. pyrrhogaster* and were also detected in the stomach contents of *Cynops ensicauda popei*, the Japanese sword-tail newt (Table S3). Because oribatid mites are known dietary sources of indolizidine alkaloids such as pumiliotoxins in poison dart frogs ^33,34^, these observations prompted us to examine whether they might also serve as a dietary source of TTX.

### Tetrodotoxin (TTX) and associated analogs in the oribatid mites

We collected oribatid mites from the soil environment of newt habitats and detected TTX using LC-MS in *Scheloribates processus* and *Galumna* sp. KM1 (Fig. 1F). LC-MS analysis revealed TTX and its analogs (4-*epi*TTX, 6-*epi*TTX, and 4,9-anhydroTTX) in both mite species, although TTX levels of several individuals were below the detection limit. Notably, the relative abundance ratios of 4-*epi*TTX and 6-*epi*TTX to TTX in mites closely matched those observed in *C. pyrrhogaster*, suggesting that oribatid mites serve as dietary sources of TTX for newts (Fig. 2A and Fig. S1). Cep-210 was detected only in *Galumna* sp., whereas trace levels of Cep-212, another putative biosynthetic intermediate of TTX ^29^, were detected in newts but not in mites (Fig. 2B). At the individual level, *Galumna* sp. (540 pg per individual) exhibited substantially higher TTX levels than *S. processus* (~20 pg per individual) (Fig. 2C). We also detected 8-deoxypumiliotoxin 193H (PTX 193H), an alkaloid previously identified in poison dart frogs and *Scheloribates* mites ^34–36^, exclusively in *S. processus* but not in *Galumna* sp. or in its predator, *C. pyrrhogaster* (Fig. S2). Certain species of poison frogs are also known to contain TTX ^13–15^, and our findings raise the possibility that mite-associated trophic pathways could contribute to toxin acquisition in other terrestrial vertebrates. Overall, we provide evidence consistent with the external (dietary) origin of TTX in newts.

**Figure 2.**
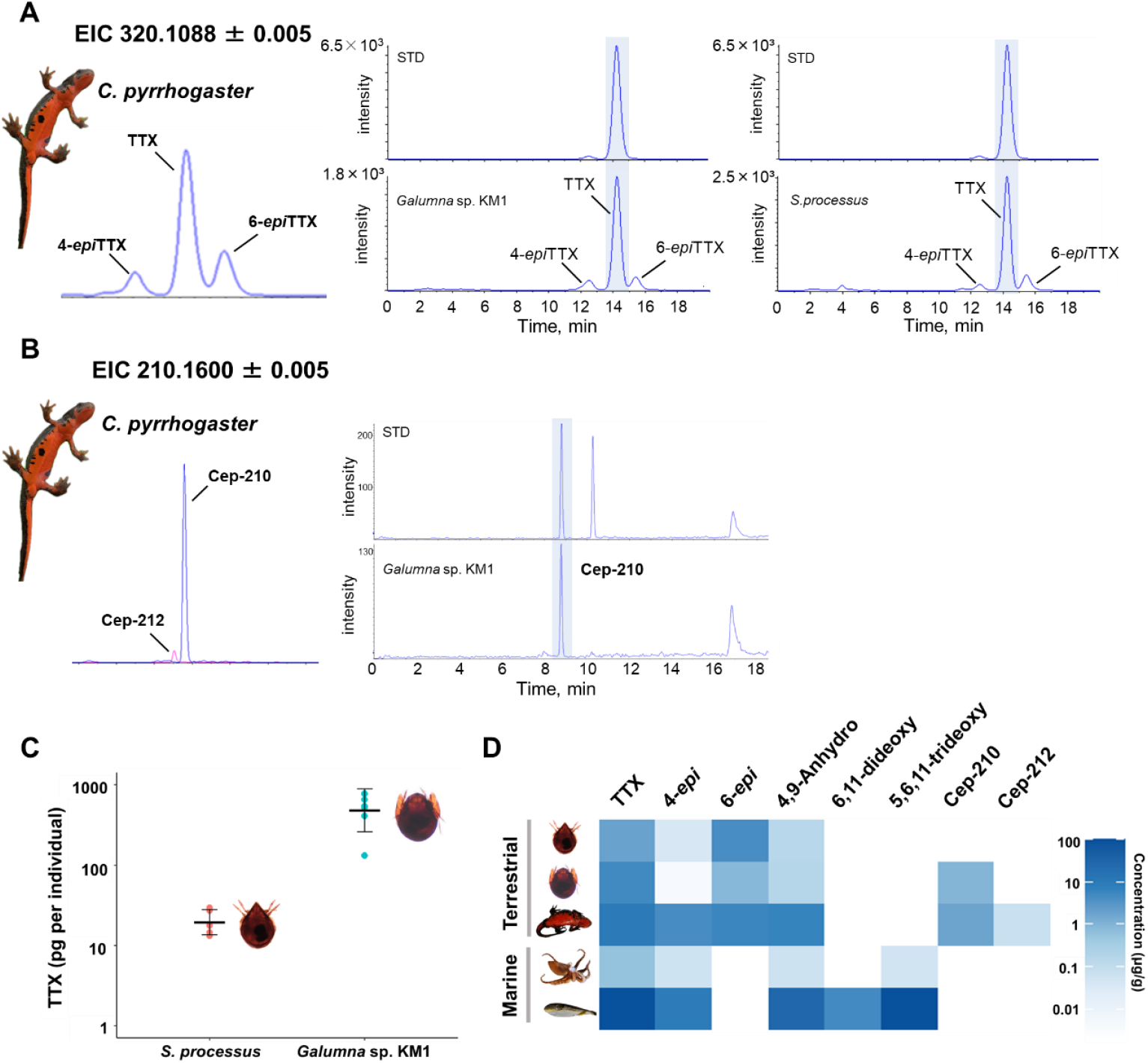
Tetrodotoxin (TTX) and associated analogs in the oribatid mites. **A**, Extracted ion chromatograms (EICs; *m*/*z* 320.1088 ± 0.005) obtained by LC-MS showing TTX and its analogs in extracts of *C. pyrrhogaster* (left), a single *Galumna* sp. KM1 individual (middle), and pooled *Scheloribates processus* (20 individuals; right). A 100 nM TTX standard is shown (STD). **B**, EICs showing Cep-210 (*m*/*z* 210.1600 ± 0.005) and Cep-212 (*m*/*z* 212.1757 ± 0.005) in *C. pyrrhogaster* (left), a single *Galumna* sp. KM1 (right). A 100 nM Cep-210 standard is shown (STD). **C**, TTX content per individual in *S. processus* and *Galumna* sp. KM1 (log scale). **D**, Heat map comparing TTX analog profiles of *S. processus* (top), *Galumna* sp. KM1 (second from top), a representative terrestrial TTX-bearing vertebrate (*C. pyrrhogaster*, middle), and representative marine taxa (blue-ringed octopus *Hapalochlaena fasciata*; pufferfish, *Takifugu poecilonotus*, from Kudo et al., 2012, 2016).

No marine-specific TTX analogs, such as TDT or 6,11-dideoxyTTX, were detected in either mite species. The TTX profiles of the mites closely resembled those of the newts, with both sharing five major analogs, TTX, 4-*epi*TTX, 6-*epi*TTX, 4,9-anhydroTTX, and Cep-210 (Fig. 2D). Experimental evidence has shown that *C. pyrrhogaster* can accumulate TTX via oral administration ^31,54^ and lacks the enzymatic capacity to metabolically interconvert TTX analogs ^54^. Together with the similarity between the TTX profiles of newts and their oribatid prey, these findings support the hypothesis that the TTX profile in newts reflects that of ingested mites rather than post-ingestion modification. Notably, although juvenile newts are capable of preying on larger, more energy-rich terrestrial arthropods (e.g., isopods such as woodlice), they nonetheless consumed large numbers of minute oribatid mites, suggesting that toxin acquisition could contribute to prey selection during the terrestrial stage. Despite the relatively small TTX content per mite, oribatid mites can account for more than 50% of the microarthropod community in soils, and their densities can reach 300,000 individuals per m^2 55^. In our feeding assays, juvenile newts consumed approximately 50 *Galumna* sp. individuals within half a day. Moreover, a single fecal pellet collected during the assays contained 35 *Galumna* sp. and 146 *S. processus* individuals. These findings suggest that mites can provide a sufficient amount of TTX to newts.

### Taxonomic Distribution of Toxin-Bearing Mites

Two TTX-bearing oribatid species, *S. processus* and *Galumna* sp., belong to the clade Brachypylina, a relatively recently derived lineage of oribatid mites ^37–39^. To assess how widely TTX occurs within Brachypylina, we collected 16 additional Brachypylina species from newt habitats and surrounding environments and further investigated whether other closely related taxa also contain TTX. These species were identified by scanning electron microscopy (SEM) and 18S rRNA gene analysis (Fig. 1F, Fig. S3, and Table S4). A molecular phylogenetic analysis was then performed using a combined dataset of 18 Brachypylina sequences in this study (including *S. processus* and *Galumna* sp.) and 59 published sequences ^38,40^ to examine the evolutionary acquisition of TTX in oribatid mites. Phylogenetic analysis revealed that *Galumna* sp. and *S. processus* are phylogenetically distant, belonging to the superfamilies Galumnoidea and Oripodoidea, respectively ^37,39^, suggesting that TTX occurrence has evolved independently in multiple oribatid lineages. Notably, these superfamilies are known to be rich sources of indolizidine alkaloids (Fig. 3 and Fig. S4) ^33,34,41,42^. However, we did not detect TTX in other species within Galumnoidea (*Galumnella nipponica* and *Trichogalumna lineata*) or Oripodoidea (*Protoribates lophothrichus* and *Peloribates longisetosus*). This pattern suggests that TTX-bearing species may not be widespread among Galumnoidea or Oripodoidea, but rather limited to certain lineages within alkaloid-bearing groups. Because the number of species analyzed in this study remains limited, the current data are insufficient to determine the evolutionary distribution of TTX in oribatid mites with confidence. Expanding chemical surveys across a broader range of taxa, particularly within Galumnoidea and Oripodoidea, will be essential for elucidating the evolutionary processes underlying the emergence and diversification of TTX association in these lineages.

**Figure 3.**
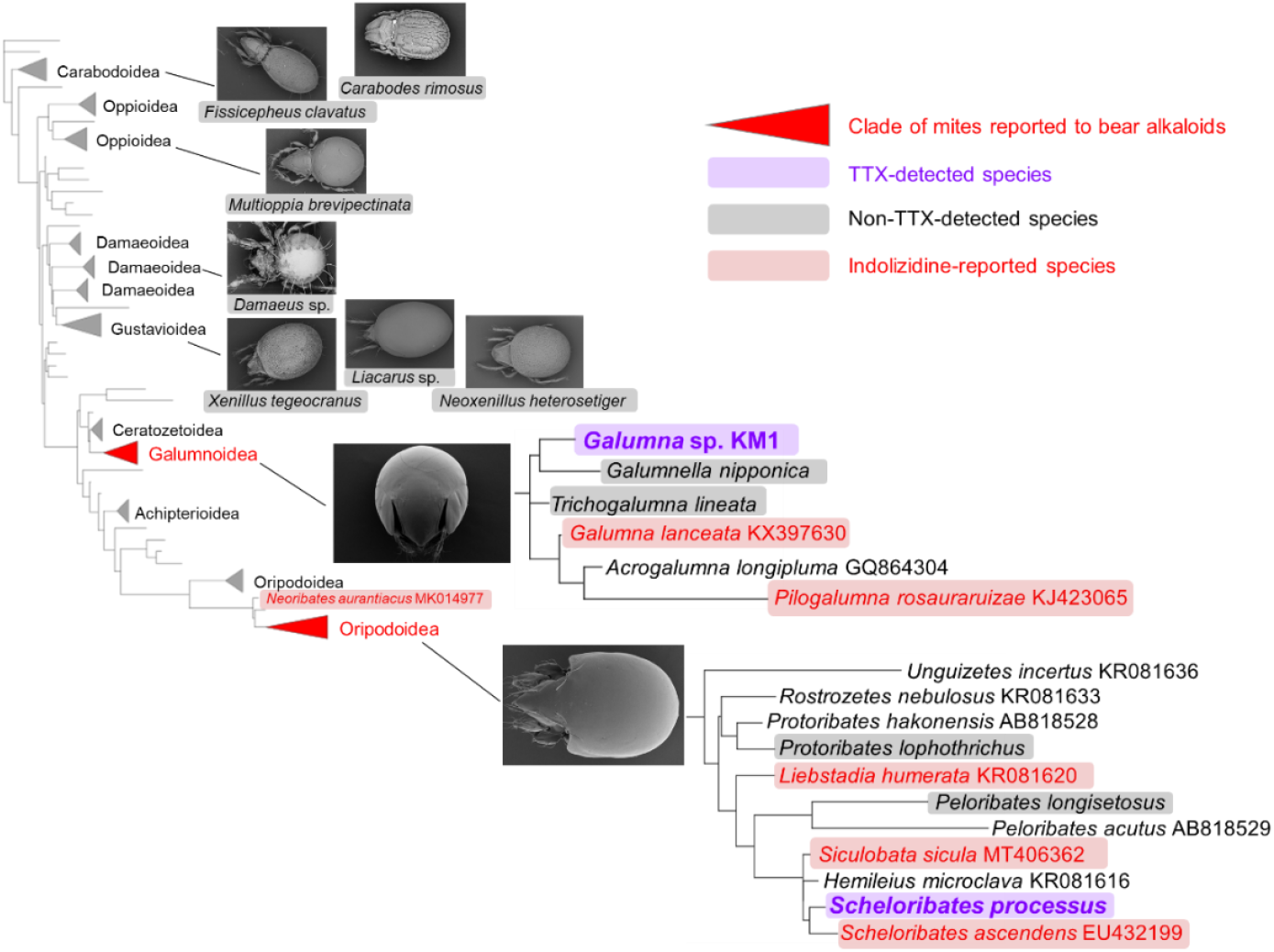
Phylogenetic relationships of the two TTX-bearing oribatid mites within Brachypylina and the distribution of reported indolizidine alkaloids. Maximum-likelihood phylogeny inferred from 18S rRNA gene sequences, including the two TTX-bearing species *Scheloribates processus* and *Galumna* sp. KM1, together with additional Brachypylina taxa sampled in this study and reference sequences from previous work. Red clades and labels denote genera or families for which indolizidine alkaloids have previously been reported in the literature (not necessarily in the sequenced species shown). Purple labels highlight the two TTX-bearing mite species, whereas gray labels indicate species in which TTX was not detected. The phylogeny is shown within the framework of Coleman and Cannatella (2022). Taxa annotated as “present in pooled sample” in that study were not treated as confirmed alkaloid-bearing here.

### TTX Localization, Potential Origin, and Ecological Function in Mites

To determine the tissue distribution of TTX, we performed immunohistochemical analyses on *Galumna* sp. and *S. processus* using an anti-TTX polyclonal antibody ^43–45^. In both species, strong immunoreactivity was observed only in the epithelial cells of the digestive tract: in *Galumna* sp., it occurred in both the caeca and ventriculus, whereas in *S. processus* it was detected only in the caeca (Fig. 4 and Fig. S5). This distribution pattern is consistent with two possibilities: (i) endogenous production by symbiotic bacteria in the digestive tissues or (ii) accumulation through the ingestion of toxic prey or microorganisms, with limited subsequent translocation to other body tissues ^50,51^, but our current data cannot discriminate between these possibilities. Several terrestrial arthropods, including *Paederus* beetles and the Asian citrus psyllid (*Diaphorina citri*), are known to harbor symbiotic bacteria that produce defensive secondary metabolites such as pederin and diaphorin, which are vertically transmitted through generations ^52,53^. The discovery of similar symbiotic bacteria in terrestrial arthropods would provide compelling evidence for a shared microbial origin of TTX across habitats. Future metagenomic and single-cell genomic approaches targeting the microbiota of oribatid mites could therefore reveal the long-sought biosynthetic gene clusters responsible for TTX synthesis.

**Figure 4.**
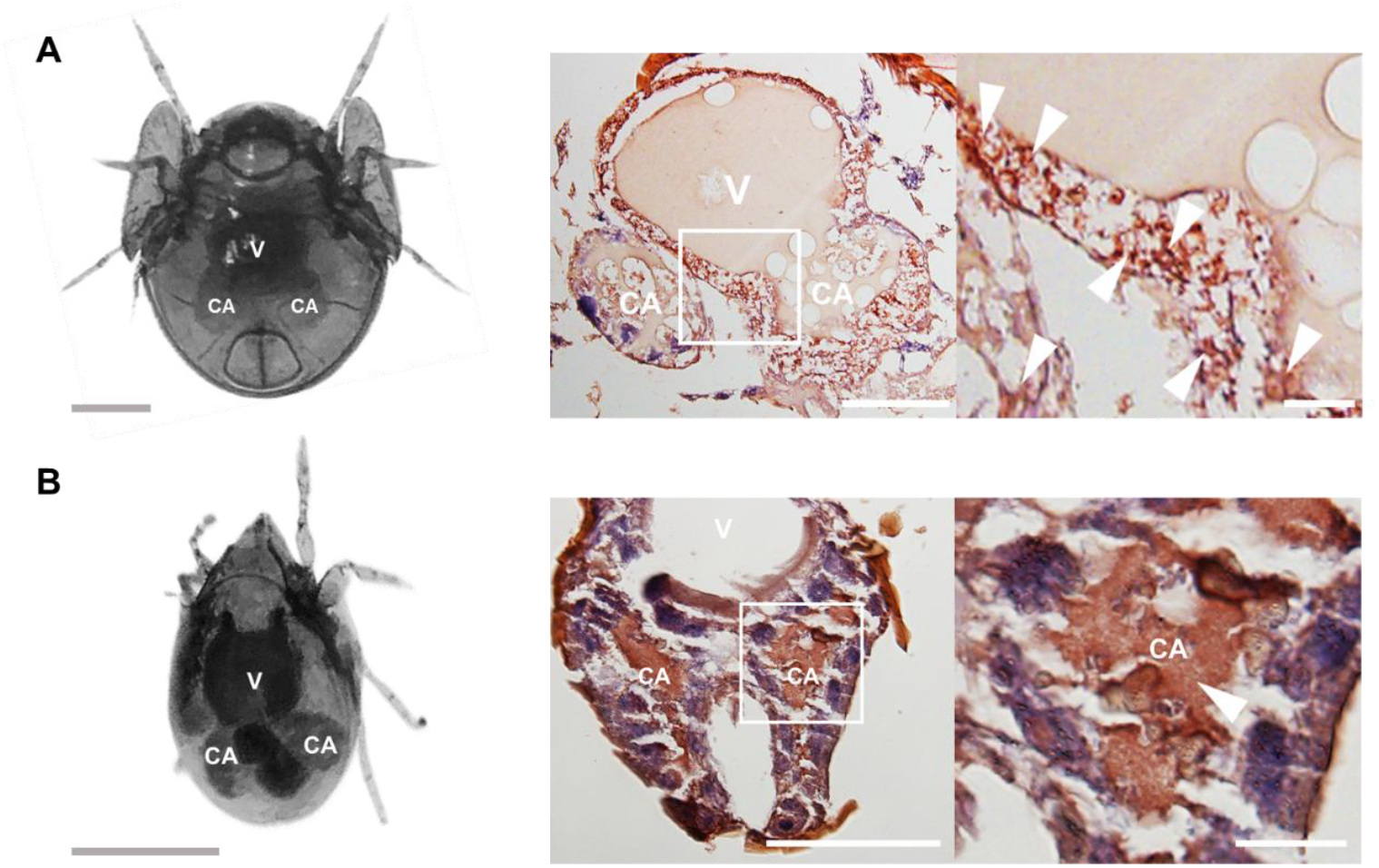
Immunohistochemical localization of TTX in *Galumna* sp. KM1 and *S. processus*. **A**, *Galumna* sp. KM1. Bright-field image of a live individual showing the digestive tract (left). Representative section stained with anti-TTX antibody showing TTX immunoreactivity (red) in the ventriculus and caeca (middle). Higher-magnification view of the boxed region in the middle panel (right). Arrowheads indicate TTX-immunoreactive epithelial cells. **B**, *S. processus*. Bright-field image of a live individual showing the digestive tract (left). Representative section stained with anti-TTX antibody (middle). Higher-magnification view of the boxed region (right). Arrowheads indicate TTX-immunoreactive epithelial cells. Scale bars: 200 µm (left panels), 100 µm (middle panels), 20 µm (right panels). V, ventriculus; CA, caeca.

The TTX content of *Galumna* sp. (0.13–0.78 ng per individual, Fig. 2C) was higher than that reported for larvae of the pufferfish *Takifugu rubripes* (0.015–0.096 ng) ^48^. Considering that *T. rubripes* larvae deter predation with a small amount of TTX, this suggests a similar defensive function in *Galumna* sp. While TTX in pufferfish larvae primarily accumulates in the liver and skin, TTX in *Galumna* sp. and *S. processus* was localized to organs associated with digestion and absorption, such as the ventriculus and caeca ^49^. This difference suggests that TTX may play distinct ecological roles in mites, and further behavioral and ecological investigations are necessary to reveal its defensive efficacy in soil ecosystems.

### Feeding Assays Using *S. processus* and *Galumna* sp

We investigated whether captive-bred, terrestrial-stage juvenile *C. pyrrhogaster* prey on TTX-bearing oribatid mites and subsequently sequester TTX in their bodies. Free-choice feeding trials showed that juveniles actively preyed on *Galumna* sp. (Fig. 5A). In a feeding assay using rhodamine-labeled *S. processus*, juveniles consumed the labeled mites, and fluorescent signals were subsequently detected in the gastrointestinal tract (Fig. 5B). We next performed a toxification experiment in which juveniles were fed TTX-enriched mites. Although the TTX content in the tail tip excised before feeding was low (0.13 ± 0.09 ng/tail tip), levels in the regenerated tail tip increased to 9.7 ± 9.0 ng/tail tip after 33 days of feeding (Fig. 5C, Table S5). In contrast, in control juveniles fed only woodlice, no increase was observed over the same period (Fig. 5C, Table S5). Notably, PTX 193H remained undetectable in the tails of juveniles fed mites after 33 days (Fig. 5D). These results demonstrate that terrestrial-stage juvenile *C. pyrrhogaster* readily consume oribatid mites and can acquire TTX through diet, resulting in measurable toxification, whereas the co-occurring alkaloid PTX 193H was not detectably transferred under these conditions.

**Figure 5.**
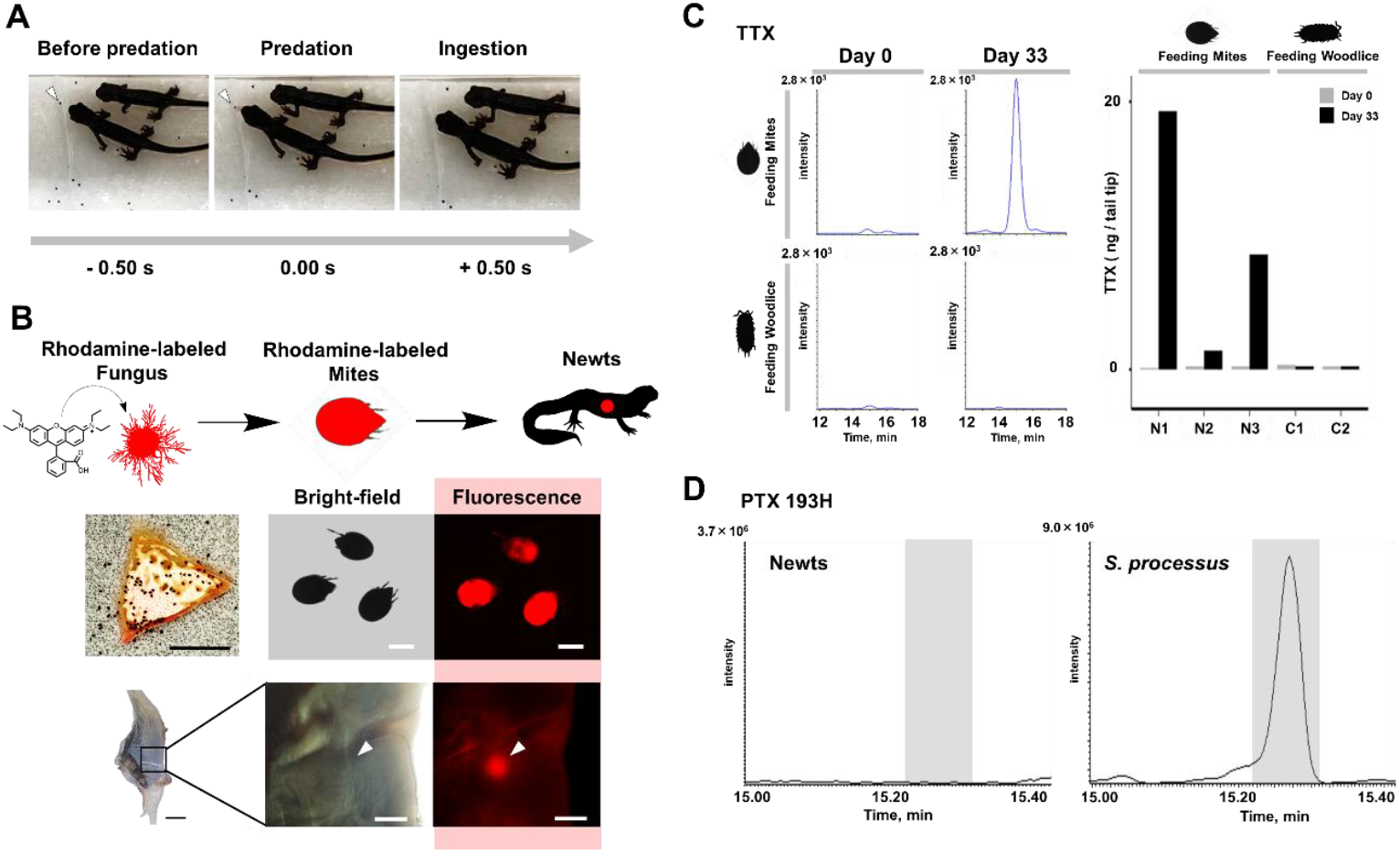
Feeding experiments with TTX-bearing oribatid mites. **A**, Free-choice feeding trials of terrestrial juvenile *C. pyrrhogaster* offered *Galumna* sp. KM1 as prey. Representative time-lapse images show sequential stages of a predation event (before predation, predation, and ingestion). Time stamps indicate time relative to the onset of predation (0 s). Negative values indicate the time before predation onset. **B**, Rhodamine-tracing feeding assay. *S. processus* were labeled with rhodamine by feeding on rhodamine-treated fungus (*Rhizoctonia candida*), and labeled mites were then offered to juvenile newts (top). Representative images show *S. processus* feeding on rhodamine-labeled fungus (middle left) and bright-field and fluorescence images of rhodamine-labeled *S. processus* (middle center and right). Bright-field images of the juvenile newt gastrointestinal tract (bottom left) and corresponding fluorescence signals from labeled mites detected in the same tissue (bottom center and right). Scale bars, 5 mm (middle left), 250 μm (middle center and right; bottom center and right), and 20 mm (bottom left). **C**, LC-MS extracted ion chromatograms (EICs) of TTX from tail-tip extracts of juvenile *Cynops pyrrhogaster*. Representative chromatograms are shown for juveniles fed TTX-bearing mites (top row) and control juveniles fed only woodlice (bottom row). For each treatment, left indicates Day 0 (before feeding) and right indicates Day 33. Bar plot shows TTX amounts (ng/tail tip) at Day 0 (gray) and Day 33 (black) for mite-fed juveniles (N1– N3) and controls (C1–C2). **D**, GC-MS ion chromatogram of an *n*-hexane extract from a tail tip of a juvenile after 33 days of feeding on mites (left) and that of *S. processus* (right). PTX 193H (gray-shaded in the TICs) was detected in *S. processus* but not in the newt tail-tip extract.

### The “Terrestrial Toxic Web” Hypothesis

The discovery of the mite *S. processus*, which harbors two distinct neurotoxins, TTX and pumiliotoxin (PTX) (Fig. 2A and Fig. S2), provides a new perspective on toxin circulation in terrestrial ecosystems. Many oribatid mites are known to feed on fungi or humus ^37^. By sequestering toxins produced by soil microorganisms or their own microbial symbionts, these mites may serve as reservoirs that facilitate toxin transfer to higher trophic levels. Our study, together with previous work, has shown that both newts and poison frogs prey on mites belonging to the families Scheloribatidae and Galumnidae ^56,57^. This raises the possibility that microbially derived toxins are transferred via mites, with TTX being delivered to newts and to frogs in genera such as *Atelopus* and *Colostethus* ^13– 15^, and PTX to other poison frogs ^36^. To date, we are not aware of evidence for overlap in toxin profiles between these amphibians. To our knowledge, PTX has not been detected in TTX-bearing Japanese fire-bellied newts (this study), and TTX has not been reported in PTX-bearing poison frogs ^15,36,58^, suggesting that distinct toxin-acquisition routes or selective retention mechanisms may operate across amphibian lineages. Therefore, convergent evolution of chemical defense acquisition mechanisms may have occurred between phylogenetically and geographically separated newts and poison frogs ^59–61^, in that both acquire defensive substances via mites. On the basis of these findings, we hereby propose the concept of a “terrestrial toxic web”, in which mites act as key intermediaries linking toxin-producing microbes with chemically defended amphibians in terrestrial ecosystems (Fig. 6). This web may represent a mode of toxin transfer that has evolved independently of marine environments. An important next step will be to test this framework by experimentally tracking toxin flow across microbial communities, mites, and vertebrate predators under controlled conditions.

**Figure 6.**
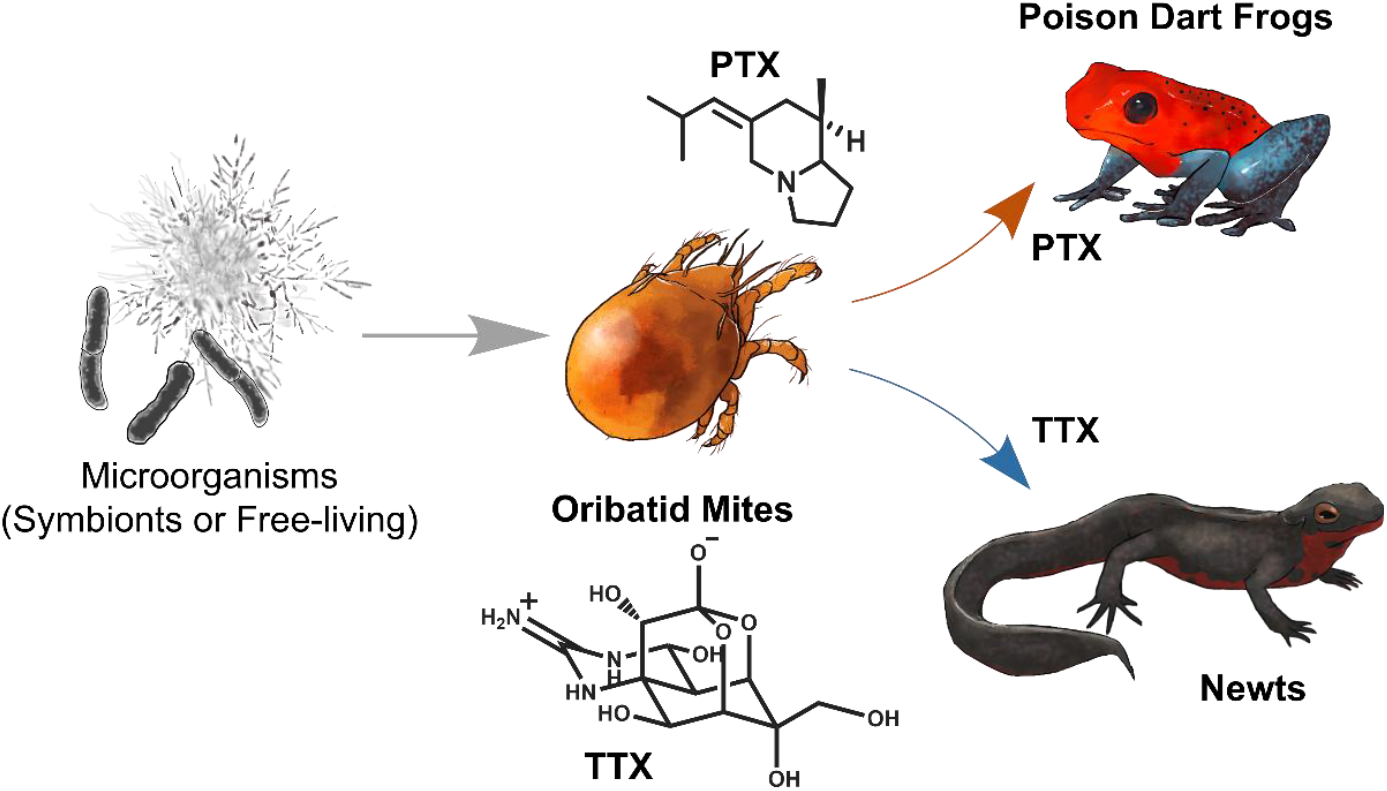
Proposed “terrestrial toxic web” hypothesis linking toxin-producing microorganisms, oribatid mites and chemically defended amphibians. Blue and brown arrows indicate toxin transfer supported by data from this study and previous studies, respectively. Gray arrows indicate hypothesized toxin transfer.

### Experimental Section

#### Collection of Mites and Newts

Individuals of *Cynops pyrrhogaster* at various developmental stages were collected in Shizuoka Prefecture, Japan, between May and November 2024. Individuals at the terrestrial stage were anesthetized and dissected immediately after capture in the field. The internal organs, including stomach contents, were removed and preserved in 95% ethanol, and the remaining body was also preserved separately in 95% ethanol. Other individuals were preserved in 95% ethanol without dissection, and the internal organs were removed in the laboratory prior to TTX analysis, with the remaining body used for TTX extraction. All samples were stored at −20 °C until analysis.

The soil-dwelling oribatid mites *Scheloribates processus* and *Galumna* sp. KM1 were collected from soil in which terrestrial-stage newts were found. A Tullgren funnel was operated for 2–4 days to collect mites from leaf litter samples. The collected mites were sorted individually and stored at −80 °C until TTX extraction. Species identification was based on both 18S rRNA gene sequencing and morphological observations: *S. processus* was identified to the species level, whereas *Galumna* sp. KM1 was identified to the genus level due to limitations in morphological resolution.

All procedures involving newts complied with institutional guidelines for the care and use of animals in research. Handling of *C. pyrrhogaster* was approved by the Animal Care and Use Committee of Kitasato University (MB2024-016).

#### Stomach Contents Analysis

Ten terrestrial-stage individuals of *C. pyrrhogaster* were used for stomach content analysis. Each individual was dissected in the field, and stomach contents were transferred to sterile tubes containing 95% ethanol and stored at −20 °C until observation. The contents were examined under a stereomicroscope, and undigested remains were identified to the lowest possible taxonomic level based on morphological characteristics.

#### Extraction and LC-MS Analysis of TTX and Cep-210

TTX and Cep-210 were extracted from oribatid mites and *C. pyrrhogaster* using 0.1% acetic acid. Crude extracts of newts were adjusted to pH 6.5–7.0 and passed through a column packed with activated charcoal. These compounds were eluted with a mixture of acetic acid, methanol, and H_2_O (2:50:48, v/v/v). LC-MS analysis was conducted using a system composed of a TripleTOF 5600+ mass spectrometer (AB SCIEX, Tokyo, Japan) coupled with an LC-20AD solvent delivery system (Shimadzu, Kyoto, Japan).

TTX analysis was performed according to the previous study ^62^. Separation was conducted using a TSK-GEL Amide-80 column (2.0 × 150 mm, 3 μm; Tosoh, Tokyo, Japan), with 16 mM ammonium formate in a mixture of H_2_O, acetonitrile, and formic acid (30:70:0.002, v/v/v) as the mobile phase. The flow rate was set to 0.2 mL/min, and the column temperature was maintained at 25 °C. ESI-MS was conducted in positive ionization mode under the following conditions: Ion Spray voltage, 4500 V; source temperature, 500 °C; nebulizer gas (GS1), 40 psi; drying gas (GS2), 70 psi; curtain gas, 25 psi. MS/MS was performed in IDA (Information Dependent Acquisition) mode, using [M + H]^+^ as the precursor ion. For the analysis of TTX in oribatid mites, *m*/*z* 320.1088 was selected as the precursor ion, and the collision energy (CE) was set to 44.0 eV.

For Cep-210 analysis, a Sun Shell C_18_ column (2.1 × 150 mm, 3.5 μm; ChromaNik Technologies, Osaka, Japan) was used. Mobile phase A consisted of H_2_O with 0.1% formic acid, and mobile phase B was acetonitrile with 0.1% formic acid. Separation was performed under the following gradient conditions (flow rate: 0.2 mL/min; column temperature: 40 °C): hold at 0% B for 2 min; linear gradient from 0% to 100% B over 20 min. ESI-MS was conducted in positive ionization mode under the following conditions: Ion Spray voltage, 5500 V; source temperature, 0 °C; nebulizer gas (GS1), 40 psi; drying gas (GS2), 70 psi; curtain gas, 25 psi.

Quantification of TTX in newt and mite samples, in addition to Cep-210 in newt samples, was performed using the external calibration method. For Cep-210, a synthetic standard was used for calibration ^63^. LC-MS analysis was carried out in positive ion mode, and quantification was performed using MultiQuant software (SCIEX, ver.3.0.2).

#### Phylogenetic Analysis

Oribatid mites used for phylogenetic analysis, excluding *S. processus* and *Galumna* sp., were collected from soil within the habitats of *C. pyrrhogaster*, as well as from forest soil in Kanagawa Prefecture, Japan. All specimens were preserved in 95% ethanol until analysis. Collected mites were identified to the lowest possible taxonomic level based on both 18S rRNA gene sequences and morphological characteristics. For DNA extraction, appendages were removed from each individual and used as the DNA source, while the remaining body was used for morphological identification and subsequently for TTX analysis.

Each appendage sample was homogenized in 3 μL of extraction buffer (0.01 M Tris-HCl pH 8.0, 0.01 M EDTA pH 8.0, 0.01% SDS, 0.1 g/L proteinase K), followed by incubation at 56 °C for 2 hours and 95 °C for 10 minutes. The resulting DNA extract was diluted 10-fold in TE buffer and used as a PCR template. The primer pair 18S_F (5′-TACCTGGTTGATCCTGCCAG-3′) and 18S_R (5′-AATGATCCTTCCGCAGGTT CAC-3′) were used to amplify the 18S rRNA gene. Amplicons were sequenced using either Sanger sequencing or the Flongle flow cell (Oxford Nanopore Technologies, UK). Sequences from 18 species of Brachypylina mites were aligned with 59 previously published 18S rRNA gene sequences used in the phylogenetic analysis by Coleman and Cannatella ^47^, which were obtained from datasets provided by Pachl et al. ^38^ and Schaefer & Caruso ^40^. Phylogenetic analysis was performed using IQ-TREE2 v2.0.7 after alignment and trimming to a uniform length. The best-fit model based on the Bayesian Information Criterion (BIC) was the SYM+G4 model, selected by ModelFinder ^64^.

Maximum-likelihood phylogenies were generated through five independent runs with 50,000 ultrafast bootstrap replicates for each. Branch support values were added to the tree with the highest log-likelihood score. The consensus tree and associated bootstrap support values are provided in the supplementary materials. The final tree was visualized using iTOL v7.

#### Immunohistochemistry

*Galumna* sp. and *S. processus* were fixed in 10% neutral buffered formalin at 4 °C for at least two nights. After fixation, samples were dehydrated through a graded ethanol series, cleared in xylene, and embedded in paraffin. Paraffin sections (8 μm thick) were prepared using a rotary microtome, deparaffinized, and blocked with 10% normal goat serum. A rabbit polyclonal anti-TTX antibody ^43–45^ was applied as the primary antibody. A biotin-conjugated anti-rabbit IgG secondary antibody and a peroxidase-conjugated streptavidin were used for detection. Immunoreactive sites were visualized using ImmPACT NovaRED (Vector Laboratories, USA), and counterstained with hematoxylin. As a negative control, antibody pre-absorbed with TTX was used and processed in parallel using the same protocol.

#### Feeding Assays Using *S. processus* and *Galumna* sp

Captive-bred juvenile *C. pyrrhogaster* were used for feeding assays. *Galumna* sp. mites were released into rearing containers of juvenile newts and then offered to them as prey. Predation events were observed and recorded under standard indoor lighting. *S. processus* were labeled with rhodamine B by feeding on rhodamine-treated *Rhizoctonia candida* (NBRC7032), using the *Rhizoctonia*-fed rearing method described previously ^65^. The labeled mites were offered to juvenile newts, and the gastrointestinal tract was examined under a fluorescence stereomicroscope.

In the toxification experiment, juveniles (n = 3) were fed mites reared on TTX-treated *R. candida*. To compensate for potential caloric deficiency during the experimental period, these juveniles were also provided with woodlice as supplemental food. As a control, additional juveniles (n = 2) were fed only woodlice for the same duration. Immediately before the feeding period, approximately 5 mm of the tail tip was excised from each individual. After 33 days of feeding, approximately 5 mm of the regenerated tail tip from the same region was excised again. The excised tail tissues were washed with distilled water, immersed in *n*-hexane, and then extracted with 0.1% acetic acid. The *n*-hexane extracts were analyzed by GC-MS for PTX detection, and the aqueous extracts were analyzed by LC-MS for TTX quantification.

## Supporting Information

Supporting Experimental Section; LC–MS/MS confirmation of TTX; GC–MS chromatograms of PTX 193H; SEM images of oribatid mites; phylogenetic tree; IHC images; detailed information on newts and their prey; information on mites used for TTX analyses and phylogenetic reconstruction; and TTX levels in tail-tip samples of juvenile newts (PDF).

### Data Availability

Nucleotide sequence data generated in this study (18S rRNA gene sequences from 18 oribatid mite taxa used for identification and phylogenetic analyses) have been deposited in the DDBJ/ENA/GenBank databases. Accession numbers are provided in Supplementary Table S4.

## Author Information

### Author Contributions

The manuscript was written through contributions of all authors. All authors have given approval to the final version of the manuscript.

### Funding Sources

This work was partially supported by JSPS KAKENHI Grant Number 22H02438 (K.T.).

### Notes

The authors declare no competing financial interests.

## Acknowledgements

We thank Dr. Masato Hirose of Kitasato University for his assistance with scanning electron microscopy techniques. We are also grateful to Mr. Mitsuru Sato for providing advice on the habitats of the Japanese fire-bellied newt. We also thank Dr. Shigeru Okada from The University of Tokyo for his assistance with GC-MS measurements.

## Notes

### Competing Interest Statement

The authors have declared no competing interest.

