## Supporting Information for "Oribatid Mites Supply Tetrodotoxin to Poisonous Newts"

**This PDF file includes:**

Supporting text
Figures S1 to S5
Tables S1 to S5
SI References

### Supplementary Experimental Section

#### Collection of *Cynops ensicauda popei* and Stomach Content Analysis

Terrestrial-stage individuals of *Cynops ensicauda popei* were collected from forests on Okinawa Island, Japan, in April 2024. Individuals were anesthetized and dissected immediately after capture in the field. Internal organs, including stomach contents, were removed and preserved in 95% ethanol and stored at −20 °C until analysis. Stomach contents from eight individuals were examined under a stereomicroscope, and undigested remains were identified to the lowest possible taxonomic level based on morphological characteristics.

#### 8-deoxypumiliotoxin 193H Analysis

Twenty individuals of *Scheloribates processus* were placed in a glass vial insert and soaked in 100 µL of *n*-hexane for 3 minutes. After removing the mites, the solvent was evaporated to dryness under reduced pressure. The residue was re-dissolved in 30 µL of *n*-hexane immediately prior to GC-MS analysis. GC-MS was conducted using a QP-2010 Ultra instrument equipped with an AOC-20 autosampler (Shimadzu, Kyoto, Japan). An InertCap-1 capillary column (0.25 mm × 60 m, film thickness 0.25 µm; GL Sciences, Tokyo, Japan) was used for separation. Helium was employed as the carrier gas at a flow rate of 1.23 mL/min. The injection volume was set to 8 µL. The oven temperature program was as follows: initial temperature of 60 °C (held for 1 min), ramped at 10 °C/min to 290 °C, and held for 10 min. Identification was based on the characteristic fragmentation pattern of 8-deoxypumiliotoxin 193H, as reported for *Scheloribates* sp. by Takada et al. [1].

#### TTX Extraction and Analysis of a Marine Organism: Blue-Lined Octopus

A female *Hapalochlaena fasciata* was obtained from an aquarium in Japan and frozen at −80 °C until TTX extraction. TTX was extracted from the whole-body tissues. Crude extracts were purified on an activated charcoal column as described for the newt samples in the main text. The purified fraction was analyzed by LC-MS under the same instrumental conditions as those used for the oribatid mite and newt samples.

### Figures

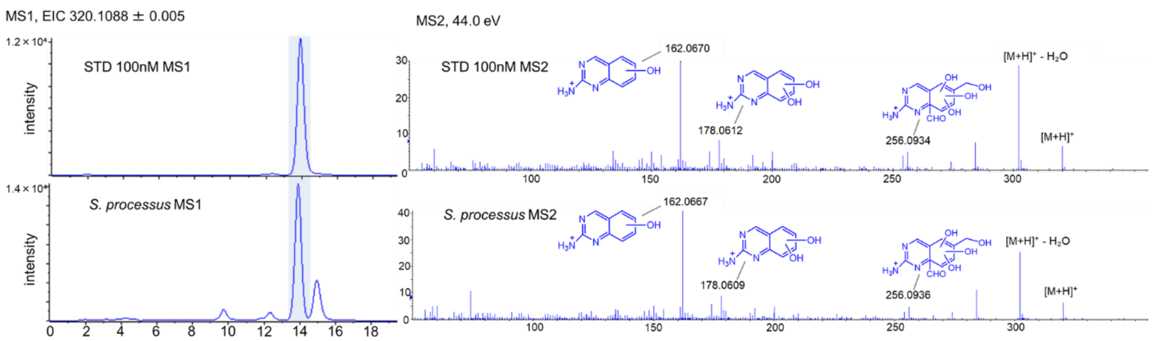


**Figure S1. LC-MS/MS confirmation of TTX in *Scheloribates processus.***

MS1 extracted ion chromatograms (EICs; *m/z* 320.1088 ± 0.005; left) and MS2 spectra (right) are shown for a 100 nM TTX standard (top) and an *S. processus* extract (bottom). TTX detected in *S. processus* showed both the same *m*/*z* and retention time in MS1, and the same MS2 fragmentation pattern, consistent with that reported by Yotsu-Yamashita et al. [2].

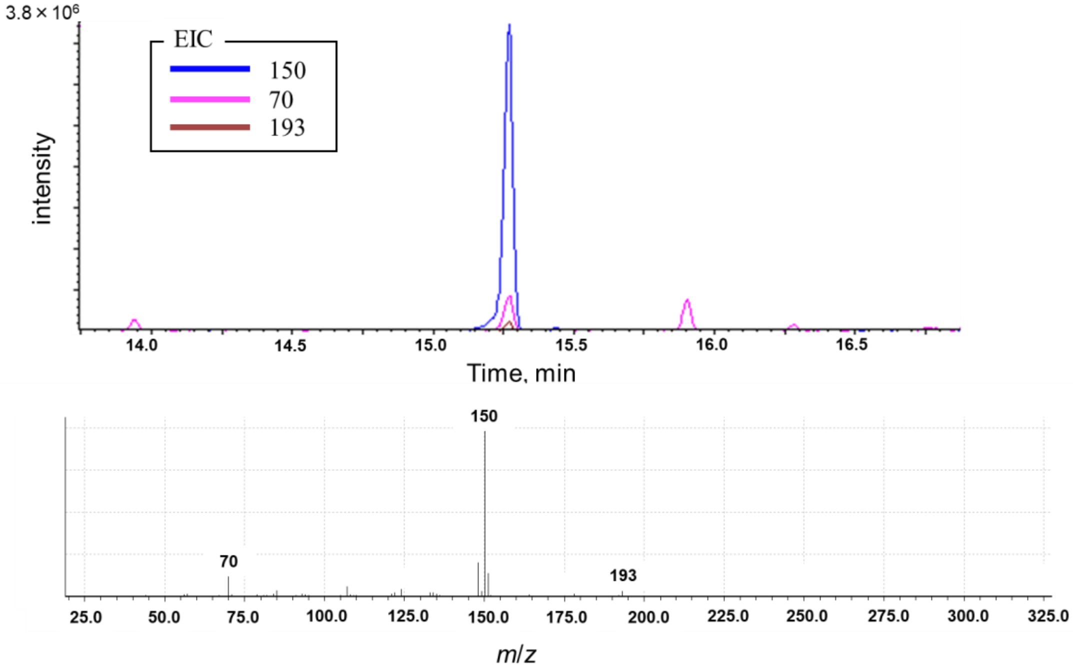


**Figure S2. GC-MS detection of 8-deoxypumiliotoxin 193H in hexane extracts of 20 *S. processus* individuals.**

Extracted ion chromatograms (top panel) monitored the molecular ion of 8-deoxypumiliotoxin 193H (*m/z* 193) and its specific fragment ions (*m/z* 70 and 150). The mass spectrum of this peak (bottom panel) was consistent with that of 8-deoxypumiliotoxin 193H previously reported from *Scheloribates* sp. [1].

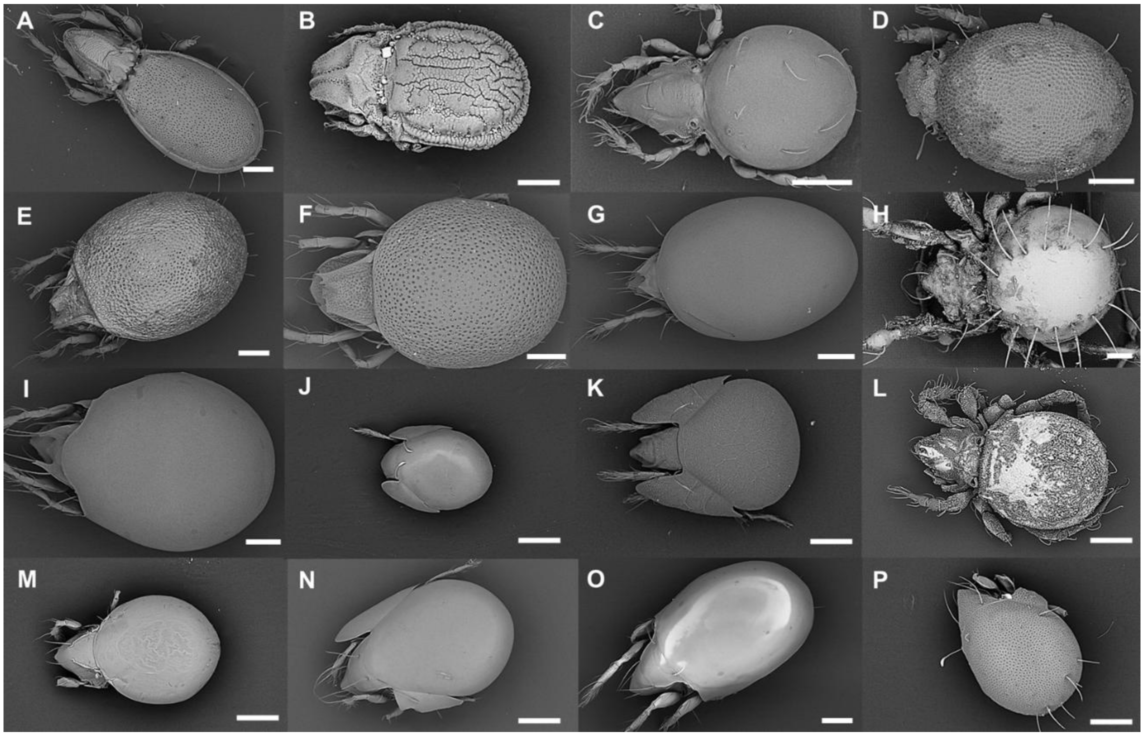


**Figure S3. Scanning electron micrographs of Brachypylina oribatid mites used for phylogenetic analysis.**

Scanning electron micrographs of 16 Brachypylina oribatid mite taxa included in the phylogenetic tree shown in Fig. 3, excluding the two TTX-bearing species *Scheloribates processus* and *Galumna* sp. KM1. Panels are labeled as follows: A, *Fissicepheus clavatus*; B, *Carabodes rimosus*; C, *Multioppia brevipectinata*; D, *Hermanniella punctulata*; E, *Xenillus tegeocranus*; F, *Neoxenillus heterosetiger*; G, *Liacarus* sp.; H, *Damaeus* sp.; I, *Euzetes* sp.; J, *Trichogalumna lineata*; K, *Galumnella nipponica*; L, *Eremobelba japonica*; M, *Oribatula sakamorii*; N, *Neoribates roubali*; O, *Protoribates lophothrichus*; P, *Peloribates longisetosus*.
Magnification, collection site, and accession numbers are provided in Supplementary Table S4. Scale bars, 100 μm.

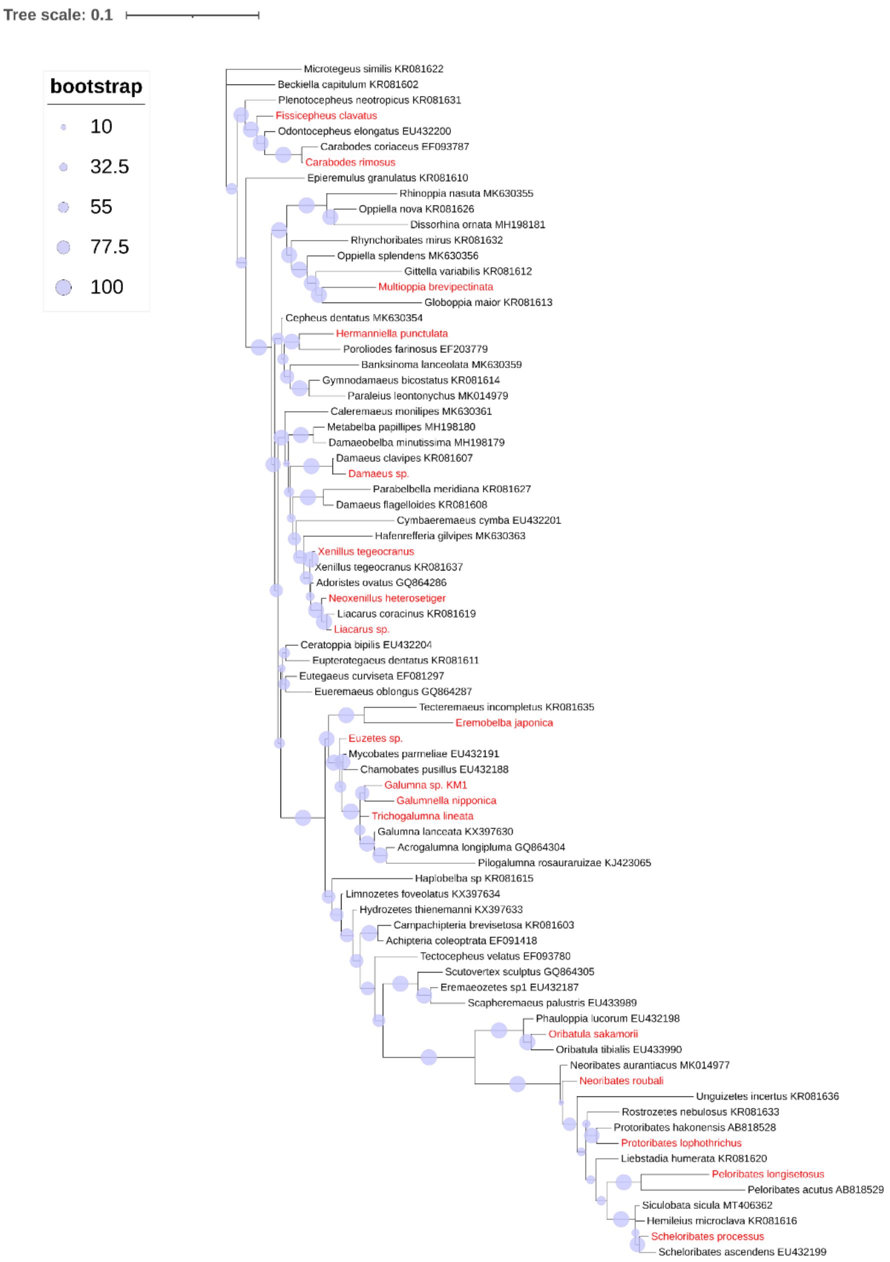


**Figure S4. Detailed phylogenetic tree of Brachypylina oribatid mites based on 18S rRNA gene sequences.**

The maximum-likelihood phylogeny was inferred from a combined dataset comprising 18 Brachypylina sequences generated in this study and 59 reference sequences from previous studies. Red labels denote taxa sequenced in this study (note that the meaning of red labels differs from that in Fig. 3). Reference sequences were obtained from the 18S rRNA datasets of Pachl et al. [3]and Schaefer and Caruso [4], which were used by Coleman and Cannatella [5] in their discussion of alkaloid acquisition during oribatid mite evolution.

A

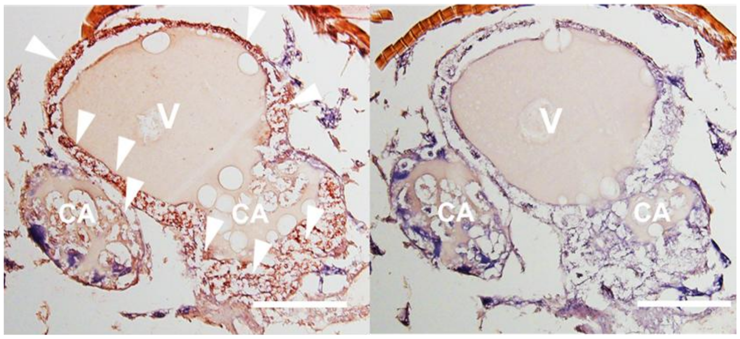


B

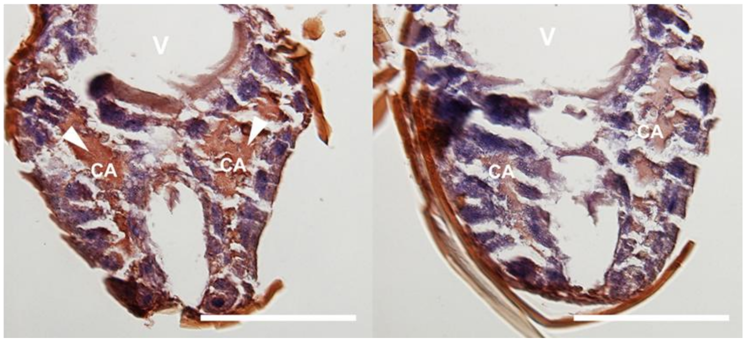


**Figure S5. Specificity control for TTX immunostaining using antigen-unabsorbed and absorbed antibodies.**

Sections corresponding to those shown in the main text (Fig. 4A and B) were stained with the unabsorbed anti-TTX antibody (left) or with the same antibody pre-absorbed with TTX (right). TTX-immunoreactive signal (red) observed with the unabsorbed antibody is indicated by arrowheads. **A**, *Galumna* sp. KM1 (same section as in Fig. 4A). **B**, *S. processus* (same section as in Fig. 4B). Scale bars, 100 µm (all panels).

**Tables**

**Table S1. Basic information of *Cynops pyrrhogaster* at each developmental stage.**

**A**, Weight, TTX amount, and Cep-210 amount in eggs and larvae. **B**, Snout–vent length (SVL), weight (after removal of internal organs), TTX amount, and Cep-210 amount in terrestrial newts. In this study, terrestrial newts with SVL less than 2.5 cm were classified as juveniles; those with SVL between 2.5 and 3.0 cm as subadults; and those larger than 3.0 cm as adults. **C**, Weight (after removal of internal organs), TTX amount, and Cep-210 amount in aquatic adults.

**A**

| Stages | Body weight (g)* | TTX (ng/individual) | Cep-210 (ng/individual) |
|---|---|---|---|
| Larva | 0.24 | ND | 18.02 |
| Larva | 0.24 | ND | 18.92 |
| Larva | 0.20 | ND | 19.97 |
| Larva | 0.11 | ND | 20.02 |
| Egg | 0.0021 | 11.71 | 1.27 |
| Egg | 0.0021 | 5.23 | 0.78 |
| Egg | 0.0025 | 2.14 | 0.98 |
| Egg | 0.0017 | 3.95 | 1.00 |
| Egg | 0.0032 | 2.90 | 0.88 |
| Egg | 0.0036 | 5.57 | 0.72 |

*In larvae, body weight refers to the weight of the newts excluding the internal organs.

**B**

| Stages* | Snout–vent length (SVL, cm) | Body weight (g)† | TTX (μg/individual) | Cep-210 (μg/individual) |
|---|---|---|---|---|
| Terrestrial-Adult | 4.0 | 1.34 | 11.59 | 0.39 |
| Terrestrial-Adult | 4.0 | 0.85 | 4.34 | 2.69 |
| Terrestrial-Adult | 4.3 | 1.68 | 16.20 | 10.78 |
| Terrestrial-Adult | 5.3 | 3.42 | 28.57 | 0.58 |
| Terrestrial-Adult | 3.8 | 1.68 | 9.93 | 0.78 |
| Terrestrial-Adult | 4.0 | 1.74 | 7.22 | 1.60 |
| Terrestrial-Adult | 4.2 | 2.07 | 8.59 | 1.13 |
| Terrestrial-Subadult | 3.0 | 1.3 | 10.55 | 1.97 |
| Terrestrial-Subadult | 3.0 | 0.95 | 10.65 | 1.00 |
| Terrestrial-Subadult | 2.9 | 0.72 | 9.13 | 1.83 |
| Terrestrial-Subadult | 2.9 | 0.77 | 7.04 | 0.65 |
| Terrestrial-Subadult | 2.9 | 0.60 | 3.34 | 0.52 |
| Terrestrial-Subadult | 2.8 | 0.76 | 4.18 | 0.25 |
| Terrestrial-Subadult | 3.0 | 0.65 | 4.50 | 0.41 |
| Terrestrial-Juvenile | 2.1 | 0.16 | 2.12 | 0.03 |
| Terrestrial-Juvenile | 2.0 | 0.16 | 2.17 | 0.03 |
| Terrestrial-Juvenile | 2.2 | 0.25 | 1.75 | 0.04 |

* For terrestrial newts: individuals with snout-vent length (SVL) between 2.5 cm and 3.0 cm are classified as Subadult, those with SVL greater than 3.0 cm are classified as Adult, and those with SVL less than 2.5 cm are classified as Juvenile.

† Body weight refers to the weight of the newts excluding the internal organs.

**C**

| Stages | Body weight (g)* | TTX (μg/individual) | Cep-210 (μg/individual) |
|---|---|---|---|
| Aquatic-Adult | 2.48 | 14.27 | 5.26 |
| Aquatic-Adult | 2.15 | 5.03 | 1.60 |
| Aquatic-Adult | 2.70 | 5.83 | 0.54 |
| Aquatic-Adult | 2.79 | 10.21 | 0.36 |
| Aquatic-Adult | 3.22 | 11.85 | 0.60 |
| Aquatic-Adult | 2.67 | 25.57 | 1.30 |
| Aquatic-Adult | 3.59 | 8.76 | 0.21 |
| Aquatic-Adult | 3.44 | 14.68 | 1.30 |

* Body weight refers to the weight of the newts excluding the internal organs.

**Table S2. Detailed information on the stomach contents of 10 terrestrial *Cynops pyrrhogaster* individuals.**

Species identification was based on morphological features and conducted to the lowest possible taxonomic level.

| Prey items | Total | Rate (%) |
|---|---|---|
| Diptera larvae | 36 | 18.7 |
| Pristomyrmex (Order: Hymenoptera, Family: Formicidae) | 1 | 0.5 |
| Strumigenys (Order: Hymenoptera, Family: Formicidae) | 1 | 0.5 |
| Monomorium (Order: Hymenoptera, Family: Formicidae) | 1 | 0.5 |
| *Eupobera* (Order: Hymenoptera, Family: Formicidae) | 1 | 0.5 |
| *Armadillidium vulgare* (Order: Isopoda, Family: Armadillidiidae) | 12 | 6.2 |
| Armadillidae (Order: Isopoda) | 2 | 1.0 |
| Isopoda (Non-conglobating) | 14 | 7.3 |
| Nematoda | 18 | 9.3 |
| *Monotarsobius* (Order: Lithobiomorpha, Family: Lithobiidae) | 2 | 1.0 |
| Chrysomelidae (Order: Coleoptera) | 1 | 0.5 |
| Curculionidae (Order: Coleoptera) | 6 | 3.1 |
| Hemiptera | 1 | 0.5 |
| Chilopoda (Unidentifiable) | 2 | 1.0 |
| Hyleoglomeris (Order: Glomerida, Family: Glomeridae) | 15 | 7.8 |
| Ampelodesmus (Order: Polydesmida, Family: Pyrgodesmidae) | 3 | 1.6 |
| Cryptocorypha (Order: Polydesmida, Family: Pyrgodesmidae) | 5 | 2.6 |
| Labidostomatidae (Order: Acari, Suborder: Prostigmata) | 1 | 0.5 |
| Prostigmata (Unidentifiable) | 3 | 1.6 |
| Gamasida (Order: Acari) | 19 | 9.8 |
| Scheloribatidae (Order: Acari, Suborder: Oribatida) | 17 | 8.8 |
| Galumnidae (Order: Acari, Suborder: Oribatida) | 7 | 3.6 |
| Peloppiidae (Order: Acari, Suborder: Oribatida) | 6 | 3.1 |
| Oribatula (Order: Acari, Suborder: Oribatida, Family: Oribatulidae) | 5 | 2.6 |
| Euphthiracaridae (Order: Acari, Suborder: Oribatida) | 1 | 0.5 |
| Camisia (Order: Acari, Suborder: Oribatida) | 1 | 0.5 |
| Limnozetes (Order: Acari, Suborder: Oribatida) | 1 | 0.5 |
| Carabodes (Order: Acari, Suborder: Oribatida) | 1 | 0.5 |
| Oribatida (Unidentifiable) | 1 | 0.5 |
| Coleoptera (Unidentifiable) | 1 | 0.5 |
| Collembola | 6 | 3.1 |
| Enicocephalidae (Order: Hemiptera) | 1 | 0.5 |
| Unidentifiable | 1 | 0.5 |
| Total | 193 | 100.0 |

**Table S3. Detailed information on the stomach contents of 8 terrestrial *Cynops ensicauda popei* individuals.**

Species identification was based on morphological features and conducted to the lowest possible taxonomic level.

| Prey items | Total | Rate (%) |
|---|---|---|
| Chrysomelidae larvae (Order: Coleoptera) | 14 | 10.8 |
| Gastropoda | 5 | 3.8 |
| Diptera larvae | 21 | 16.2 |
| Discolomatidae (Order: Coleoptera) | 9 | 6.9 |
| Cryptocorypha (Order: Polydesmida, Family: Pyrgodesmidae) | 17 | 13.1 |
| Hyleoglomeris (Order: Glomerida, Family: Glomeridae) | 3 | 2.3 |
| Rhinotus (Order: Polyzoniida, Family: Siphonotidae) | 1 | 0.8 |
| Diplopoda (Unidentifiable) | 6 | 4.6 |
| Collembola | 8 | 6.2 |
| Gamasida (Order: Acari) | 8 | 6.2 |
| Oribatida (Order: Acari) | 11 | 8.5 |
| Astigmata (Order: Acari) | 1 | 0.8 |
| Prostigmata (Order: Acari) | 1 | 0.8 |
| Amphipoda (Family: Talitridae) | 4 | 3.1 |
| Strumigenys (Order: Hymenoptera, Family: Formicidae) | 4 | 3.1 |
| Carebara (Order: Hymenoptera, Family: Formicidae) | 2 | 1.5 |
| Monomorium (Order: Hymenoptera, Family: Formicidae) | 2 | 1.5 |
| Pyramica (Order: Hymenoptera, Family: Formicidae) | 1 | 0.8 |
| Pristomyrmex (Order: Hymenoptera, Family: Formicidae) | 1 | 0.8 |
| Formicidae (Unidentifiable) | 1 | 0.8 |
| Isopoda | 2 | 1.5 |
| Pselaphinae (Order: Coleoptera, Family: Staphylinidae) | 2 | 1.5 |
| Heteroptera (Order: Hemiptera) | 1 | 0.8 |
| Araneae | 2 | 1.5 |
| Coleoptera (Unidentifiable) | 2 | 1.5 |
| Hymenoptera (Unidentifiable) | 1 | 0.8 |
| Total | 130 | 100.0 |

**Table S4. Information on Brachypylina oribatid mites used in the phylogenetic tree shown in Fig. 3.**

| Taxon | Collection site | SEM magnification | TTX detected | Analogs detected | Accession No. |
|---|---|---|---|---|---|
| *Fissicepheus clavatus* | Shizuoka | 350× | × | — | PX698614 |
| *Carabodes rimosus* | Kanagawa | 550× | × | — | PX698613 |
| *Multioppia brevipectinata* | Shizuoka | 660× | × | — | PX698606 |
| *Hermanniella punctulata* | Shizuoka | 490× | × | — | PX698608 |
| *Xenillus tegeocranus* | Shizuoka | 370× | × | — | PX698610 |
| *Neoxenillus heterosetiger* | Shizuoka | 340× | × | — | PX698611 |
| *Liacarus* sp. | Shizuoka | 360× | × | — | PX698612 |
| *Damaeus* sp. | Shizuoka | 370× | × | — | PX698609 |
| *Euzetes* sp. | Shizuoka | 360× | × | — | PX698616 |
| *Trichogalumna lineata* | Shizuoka | 530× | × | — | PX698617 |
| *Galumna* sp. KM1 | Shizuoka | 100× | ● | 6-*epi* TTX, 4-*epi* TTX, 4,9-anhydroTTX, Cep-210 | PX698618 |
| *Galumnella nipponica* | Kanagawa | 450× | × | — | PX698615 |
| *Eremobelba japonica* | Shizuoka | 680× | × | — | PX698607 |
| *Oribatula sakamorii* | Shizuoka | 740× | × | — | PX698619 |
| *Neoribates roubali* | Shizuoka | 540× | × | — | PX698622 |
| *Protoribates lophothrichus* | Shizuoka | 510× | × | — | PX698623 |
| *Scheloribates processus* | Shizuoka | 240× | ● | 6-*epi* TTX, 4-*epi* TTX, 4,9-anhydroTTX | PX698621 |
| *Peloribates longisetosus* | Shizuoka | 540× | × | — | PX698620 |

Symbols: **●, detected; ×, not detected.**

**Table S5. TTX levels in tail-tip samples of juvenile newts before and after feeding experiments.**

TTX amounts (ng/tail tip) and concentrations (ng/g) were quantified at Day 0 (before feeding) and Day 33 (after 33 days). Newt1–3 were fed TTX-bearing mites, whereas CTRL1–2 were fed only woodlice.

| | Day 0 (TTX, ng/tail tip) | Day 0 (TTX, ng/g) | Day 33 (TTX, ng/tail tip) | Day 33 (TTX, ng/g) |
|---|---|---|---|---|
| Newt 1 | 0.026 | 32 | 19.26 | 7131 |
| Newt 2 | 0.19 | 105 | 1.32 | 294 |
| Newt 3 | 0.19 | 132 | 8.57 | 3172 |
| CTRL 1 | 0.26 | 650 | 0.22 | 555 |
| CTRL 2 | 0.21 | 268 | 0.16 | 205 |
